# Immunogenicity of COVID-eVax Is Moderately Impacted by Temperature and Molecular Isoforms

**DOI:** 10.1101/2022.12.29.522202

**Authors:** Federico D’Alessio, Lucia Lione, Erika Salvatori, Federica Bucci, Alessia Muzi, Giuseppe Roscilli, Antonella Conforti, Mirco Compagnone, Eleonora Pinto, Gianfranco Battistuzzi, Luigi Aurisicchio, Fabio Palombo

## Abstract

DNA integrity is a key issue in gene therapy and genetic vaccine approaches based on plasmid DNA. In contrast to messenger RNA that requires a controlled cold chain for efficacy, DNA molecules are considered to be more stable. In this study, we challenged this concept by characterizing the immunological response induced by a plasmid DNA vaccine delivered using electroporation. As a model, we used COVID-eVax, which is a plasmid DNA vaccine that targets the receptor binding domain (RBD) of the SARS-CoV-2 spike protein. Increased nicked DNA was produced by using either an accelerated stability protocol or a lyophilization protocol. Surprisingly, the immune response induced in vivo was only minimally affected by the percentage of open circular DNA. This result suggests that plasmid DNA vaccines, such as COVID-eVax that has completed a phase I clinical trial, retain their efficacy upon storage at higher temperatures and this feature may facilitate their use in low-/middle-income countries.

## 1. Introduction

Non-viral vaccines against SARS-CoV-2 reached global diffusion with the COVID-19 pandemic. Significant success in preventing infection and deaths was achieved in a fast unprecedented timeframe due to favorable characteristics such as rapid turnaround production, low toxicity, and high immunogenicity. These features are now recognized as the key parameters for pandemic preparedness. This class of vaccines includes messenger RNA (mRNA)-based vaccines and plasmid DNA vaccines delivered in different modalities. Plasmid DNA as a drug must be highly purified to remove all residual bacterial proteins and the host DNA [1]. In addition, for extended stability at a convenient storage temperature, contaminating nucleases must be removed. Among its distinctive advantages, plasmid DNA readily and quickly renatures under many conditions with no loss of biological activity. Differently from protein precipitation or aggregation, plasmid DNA usually requires chemical modification to produce an irreversible loss of biological activity, thus, indicating DNA chemical integrity as the primary endpoint of structural stability studies. In principle, this offers the opportunity to replace measurements of biological activity with high-resolution chemical analysis and, therefore, to evaluate plasmid DNA-based products similarly to small molecule drugs rather than biological entities.

Monitoring the stability of plasmid DNA over long periods is mainly dependent on the chemical integrity of the phosphodiester backbone. The introduction of a single break in the DNA backbone converts supercoiled (SC) plasmid DNA to open circular (OC) plasmid DNA, or a double nick on both strands leads to linear (L) DNA. Monitoring the conversion of SC to OC DNA provides a convenient and sensitive assay to assess DNA stability. There are different methods to monitor the conversion of SC plasmid DNA to the OC and L forms of DNA, such as agarose gel electrophoresis and various chromatographic techniques. These two methods are included in the release tests of injectable plasmid DNA [1].

DNA delivery methods are key to achieving optimal gene expression and immunogenicity. Recently, we have completed a phase I clinical trial with COVID-eVax (NCT04788459), a naked plasmid DNA expressing the receptor binding domain (RBD) of the SARS-COV-2 spike protein, delivered by electroporation (EP). COVID-eVax has shown potent immunologic efficacy and protection in preclinical models of infections [2,3]. In addition to the DNA-EP delivery [4,5], naked DNA has been historically administered as such [6], by air pressure or complexed with different carriers [7]. One of the advantages of DNA is its stability, which is a function of the buffer composition and storage temperatures [8]. Nicked DNA increases as a function of the temperature, requiring a cold chain to maintain DNA integrity. According to Pharmacopeia, SC plasmid DNA in injectable DNA must be higher than 80% [1]. However, the impact of nicked DNA on biological functions such as gene expression or as an immune response in vivo was not evaluated in the context of electroporation delivery. In this study, we characterize the impact of the nicked DNA produced in an accelerated stability test [9] or lyophilization on COVID-eVax expression levels by performing an in vitro potency assay and evaluating the immune response induced in a mouse model upon intramuscular delivery by EP. Our immunological data suggest that the EP delivery of up to 50% relaxed DNA does not affect DNA-based vaccine immunological efficacy.

## 2. Materials and Methods

### Plasmid DNA Manufacturing

The production of the engineering run of the plasmid vector COVID-eVax was carried out at Biomay (Vienna Austria) under current principles of good manufacturing practice (GMP) and following the guidelines for the “Manufacture of biologically active substances and medicinal products for human use” (Annex 2 of the EU-GMP guideline) and “Manufacture of investigational medicinal products” (Annex 13 of the EU-GMP guideline). The final concentration was adjusted to 4 ± 0.4 g/L product. The DNA was stored in 2 ml glass tubes at -20 °C and used for all experiments. The DNA was stored in PBS, pH of 7.3, at -20 °C, and tested for stability for 2 years.

### Freeze-drying

The process was carried out using a VirTis SP Scientific Advantage EL-85 Freeze Dryer. Samples of COVID-eVax (500 µL) in 1.5 mL glass vials were allowed to freeze at -50 °C for 5 hours under a light vacuum (about 500 mbar), and then the following conditions were applied:

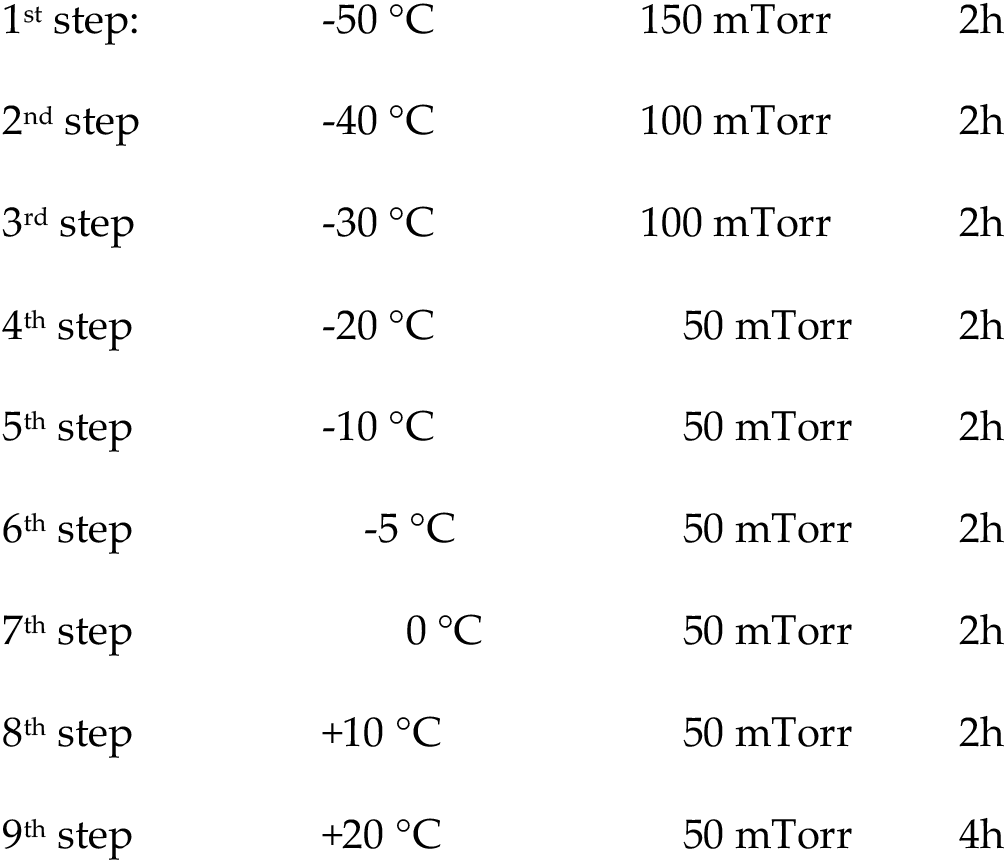

At the end of the sequence, the vacuum was broken by fluxing ultra-pure nitrogen, and vials were quickly capped under an inert atmosphere.

### Analytical Methods

The plasmid vector COVID-eVax that encodes for the receptor binding domain (RBD) of the spike protein of the coronavirus Sars-CoV-2 was produced following a GMP-like protocol, at Biomay (Austria), stored in 2 ml glass tubes at -20 °C, 4 ug/uL, and used for all experiments. The molecular weight and the supercoiled/open coiled ratio of the pDNA were analyzed by running 200 ng of plasmid DNA on each lane on a 1% agarose TBE gel stained with SYBR-Safe DNA (Thermo Fisher Scientific,USA) for 2 h at 45 V. The gel images were acquired using a ChemiDoc™ MP Imaging System and quantification of the pDNA isoform ratios were determined through densitometric analysis using the software ImageLab 6.0.1 (BioRad, USA).

To set up the HPLC analytical method, reference standards were generated by enzymatic restriction, OC was generated with Nt.BbvCI (New England Biolabs, USA) and L DNA was generated with HindIII-HF (New England Biolabs, USA). A CIMac pDNA 0.3 mL weak anion-exchange analytical column (Sartorius, Germany) was used with a Shimadzu

Prominence HPLC system (Shimadzu, Japan) equipped with a SIL-20ACHT UFLC autosampler, a column heater set to 25 °C, and an SPD-20AV detector. Absorbance was monitored at 260 nm and the Shimadzu LabSolutions software was used for peak integration to obtain area and area %. The equilibration buffer (A) was 200 mM Tris–HCl with pH 8.0; the elution buffer (B) was 200 mM Tris–HCl, 1 M NaCl, with pH 8.0. All buffers were filtered through 0.2 µm filters before use. The flow rate was 1.0 mL/min throughout. Each sample was injected in a 10 µL volume containing 1 µg supercoiled DNA, 0.2 µg linear DNA, or 0.2 µg open circular DNA. A linear gradient was developed from 65% to 85% buffer B in 15 min.

### Potency Assay

The HEK293 cells were transfected using the same amount of each DNA sample, according to the Lipofectamine 2000 manufacturer’s instructions (ThermoFisher Scientific,USA). Then, 72 h later, supernatants were collected and evaluated by Sandwich Elisa assay. Briefly, Maxisorp 96-well plates (Nunc) were coated with 1 ug/ml of the anti-RBD antibody 5B7 (generated in Takis) in PBS1X overnight at 4 °C, and after a wash in PBS1X-Tween 0.05% (PBST), they were blocked with 3% BSA-PBST 1 h at RT in agitation. Scalar doses of each supernatant in 1% BSA-PBST were added, and purified RBD [3] was used as the reference standard control. After o/n incubation at 4 °C, plates were washed and the SARS-CoV spike antibody, Rabbit PAb, Antigen Affinity Purified (Sino biological Cat: 40150-T62-COV2 100) was added and incubated for 3 h at RT. Finally, plates were washed and incubated with the Goat Anti-Rabbit IgG (H + L)-HRP Conjugate (Biorad, #1706515) diluted 1:2000 in BSA 1%/PBST, 1 h at RT, and developed by adding 50 ul/well of TMB (3,3’ s,5,5’ tetramethylbenzidine) liquid substrate (Sigma, Germany). Absorbance was measured at 450 nm using the ELISA plate reader Tecan. Data were plotted and analyzed using GraphPad to quantify RBD concentration in the supernatant of transfected HEK293 cells.

### Analysis of Immune Responses

The in vivo experimental procedures were all approved by the local animal ethics council and the ethical committee of the Italian Ministry of Health, authorization # 586/2019-PR. The mouse experiments were conducted with 6-week-old C57BL/6 female mice (Envigo. USA), as described in the figure legends. Mice were vaccinated with 10 ugs of plasmid DNA delivered by electroporation (DNA-EP) in the quadriceps, following a prime-boost vaccination regimen (Days 0 and 28). DNA-EP was performed using a Cliniporator device (Igea, Italy), using a needle electrode (electrode N-10-4B). At different timepoints, antibody and cell-mediated immune responses were analyzed.

### ELISA Assay

The ELISA assay was performed as previously reported [3]. Briefly, the plates were functionalized by coating the RBD-6xHis protein and blocked with 3% BSA-0.05% Tween 20-PBS. Mouse sera were added at a dilution of 1/300 in 1% BSA-0.05% Tween 20-PBS and diluted 1:3 up to 1/218700, in duplicate. After incubation overnight at 4 °C, antibody levels were detected using a secondary anti-murine IgG conjugated with alkaline phosphatase and the substrate for alkaline phosphatase yellow (pNPP) liquid substrate system for ELISA (cat. P7998, Sigma). After 30 minutes of incubation, the absorbance at 405 nm was measured using an ELISA reader. IgG antibody titers against the RBD protein were evaluated at two timepoints (Days 28 and 35). Endpoint titers were calculated by plotting the log10 OD against the log10 sample dilution. A regression analysis of the linear part of the curve allowed the calculation of the endpoint titer. An OD of 0.2 was used as a threshold.

### ELISpot Assay

The assay was performed on splenocytes from vaccinated and control mice, according to the manufacturer’s instructions (Mabtech, Sweden) as previously described [4]. Splenocytes were plated at 4 × 10^5^ and 2 × 10^5^ cells/well, in duplicate, and stimulated with a pool of RBD peptides at a final concentration of 1 μg/ml. The next day, plates were developed according to the kit manufacturer’s instructions. The results were spot-forming cells (SFC) per million splenocytes counted with an automated ELISPOT reader (Aelvis ELIspot reader, A.EL.VIS Gmbh).

## 3. Results

### 3.1. COVID-eVax Profile Changes as a Function of Storage Temperature and Lyophilization

To establish a stability profile, the plasmid COVID-eVax was placed in stable storage at a standard temperature of - 20 °C. The DNA was analyzed at 0, 6, 12, 18, and 24 months for gene expression and compared with the engineering run as a reference. A potency assay for the evaluation of the protein expressed by the plasmid was established and validated (see Materials and Methods). COVID-eVax encodes for the RBD fused to the TPA leader sequence, which drives the protein in the supernatant of transfected HEK293 cells. As shown in Figure 1A, RBD produced at each timepoint matched the stability criteria, which was a variation of expression less than two standard deviations.

**Figure 1.**
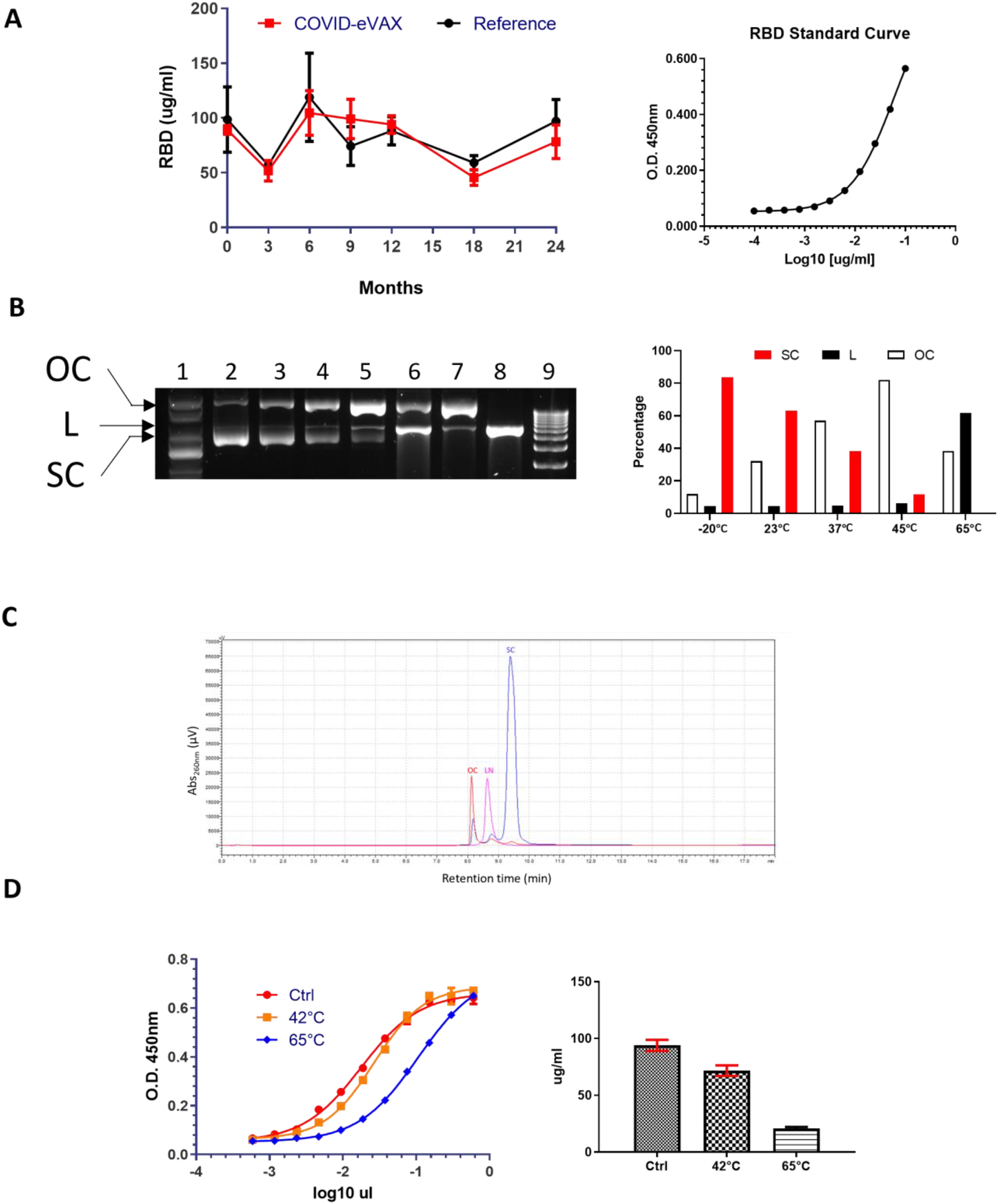
In vitro characterization of accelerated DNA stability: (**a**) Potency assay with GMP COVID-eVax with respect to reference engineering run DNA (reference) overtime and standard curve with purified RBD; (**b**) agarose gel electrophoresis of COVID-eVax stored at different temperatures, lanes are as follows: (1) supercoil marker DNA, (2) DNA stored at -20 °C, (3) DNA kept at 23 °C, (4) DNA kept at 37 °C, (5) DNA kept at 45 °C, (6) DNA kept at 65 °C, (7) control nicked DNA, (8) control linearized DNA, (9) 1 kb linear DNA ladder, and on the right side of the panel, quantification of the different DNA forms at selected temperatures as described in Materials and Method; (**c**) Representative overlayed chromatograms of LN, OC, and SC of COVID-eVax; (**d**) Potency assay of two representative temperatures (42 °C and 65 °C), standard curve (Ctrl), and quantification are shown in the graph.

Having established that COVID-eVax is stable at -20 °C for at least 2 years without losing its physicochemical and gene expression properties, we asked what is the impact of OC DNA on the immune responses induced by DNA-EP.

To this end, we adopted the strategy used for an accelerated stability test [9]. The COVID-eVax plasmid was incubated at different temperatures for 20 days, and then DNA integrity was analyzed by gel electrophoresis (Figure 1B). As expected, the percentage of OC DNA increased as a function of temperature, depicted by the gel electrophoresis and confirmed by HPLC analysis (Figure 1C). More specifically, the SC form showed a linear decrease from 84% to 12% in the DNA kept at 45 °C, while the OC form, as expected, increased in the opposite direction from 12% to 82%. The percentage of linear DNA was stable at around 5% up to 45 °C but increased dramatically to 62% at 65 °C. Then, we looked at a biological function in vitro using the potency assay previously described (Figure 1D). A reduced gene expression was observed only with the highly degraded DNA kept at 65 °C. These results suggest that plasmid DNA with an increased percentage of OC DNA, such as that observed for the sample incubated at 45 °C, maintains the expression levels and biological functions.

### 3.2. COVID-eVax Immune Response Is Poorly Impacted by Molecular Isoform Composition

To evaluate the impact of DNA integrity on the immune response induced by electroporation delivery, C57/B6 mice were vaccinated with COVID-eVax DNA treated as reported in Figure 2. This vaccination protocol was published recently with a prime injection followed by a boost at Day 28 [3]. One week after the boost, spike-specific T-cell responses were evaluated in the spleen upon overnight stimulation with the peptide pool covering the entire RBD sequence of the spike protein. Significant immune responses were observed in 60% (three out of five) of mice vaccinated with DNA kept at 23 °C, and in 80% (four out of five) of mice vaccinated with DNA kept at 45 °C (Figure 2A).

**Figure 2.**
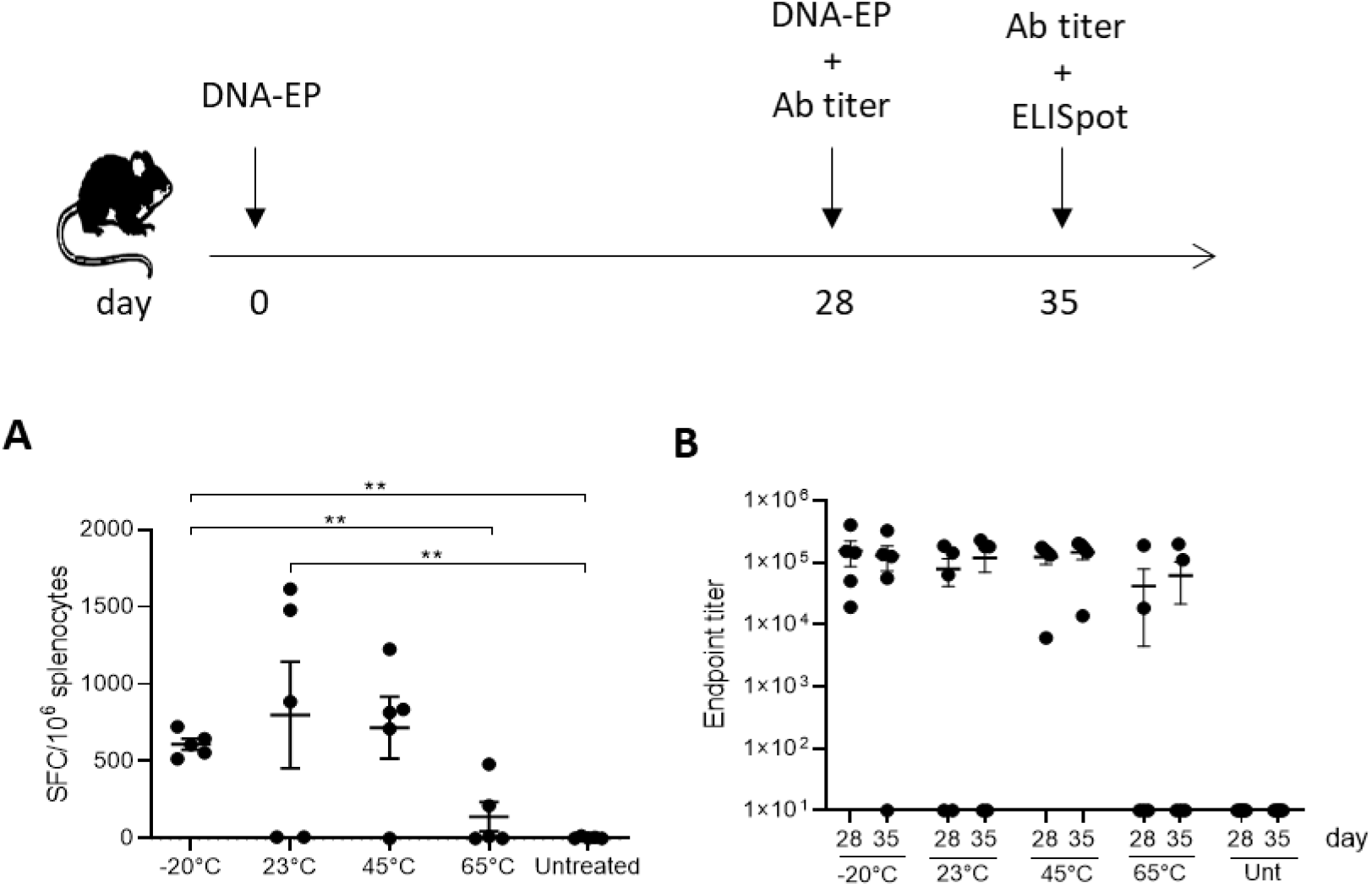
OC COVID-eVax vaccine delivered by EP is immunogenic. Groups of C57Bl/6 mice (*n* = 5) were vaccinated on Days 0 and 28 and sacrificed on Day 35 for IFN-γ ELISpot analysis. Sera were collected on Days 28 and 35 for binding antibodies titer analysis: (**a**) IFNγ-producing T cells measured by ELISpot assay on splenocytes restimulated with RBD peptide pool; (**b**) total RBD-specific IgG endpoint titers measured by ELISA assay in sera collected on Days 28 and 35. The data are both from one of two experiments; each symbol represents an individual sample with the error bars representing the SEM.

No statistical difference was observed with respect to immune responses induced by reference DNA (−20 °C). Parallel results were observed with the antibody response measured before the boost vaccination or after one week. The average titer of mice vaccinated with DNA kept at 23 °C and 45 °C was comparable with that of DNA stored in the reference condition (−20 °C).

To further correlate the in vitro and in vivo biological functions with the percentage of OC DNA, we generated nicked DNA using a different protocol. The DNA formulated in PBS was dried out and kept as a powder for two weeks at room temperature. Then, the DNA was resuspended in injectable water and analyzed on an agarose gel for integrity. The results showed that in this condition the SC, OC, and L isoforms were 56%, 44%, and 0%, respectively (Figure 3).

**Figure 3.**
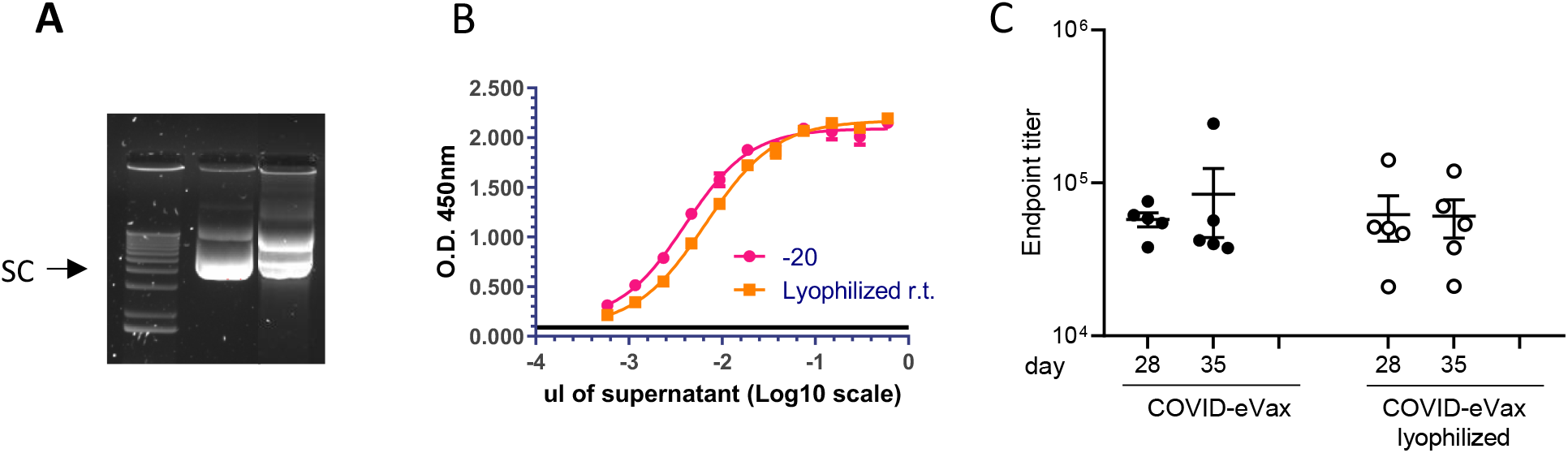
COVID-eVax elicited a similar antibody response in the lyophilized form: (**a**) DNA was lyophilized to induce DNA nicking (Materials and Methods); (**b**) potency assay showing the gene expression of DNA in panel A; (**c**) endpoint titers measured by ELISA assay of C57Bl/6 mice (*n* = 5) vaccinated via the DNA-EP depicted in Figure 1. Each symbol represents an individual sample with the error bars representing the SEM.

The potency assay showed a loss of expression by 38%. However, the antibody titer induced in mice vaccinated with the same immunization schedule depicted in Figure 2 showed similar antibody titers as reference DNA (Figure 3). Taken together, these results show that the OC DNA, up to 82%, did not affect either the in vitro expression or the induction of immune response in vivo.

## 4. Discussion

While the pandemic has accelerated the development of new vaccine technologies, it has also shown the importance of making these vaccines equitably accessible to the people who need them. One of the factors that complicate the worldwide distribution and administration of COVID-19 vaccines is their need for a cold chain for shipment and storage. Although this issue has been faced in the smallpox global vaccination campaign [10], it became more evident for the novel mRNA-based vaccines that currently require continuous storage at ultra-cold temperatures, which was not feasible in many parts of the world. The need for thermostable vaccines had already been recognized before the COVID-19 pandemic, for example, in a study by the Vaccine Innovation Prioritization Strategy (VIPS), in which the ability to withstand heat exposure was identified as the most desired characteristic for vaccines used in outreach and campaign settings by experienced immunization staff. Lack of vaccine thermostability leads to limited or no access to vaccinations by people living in remote areas or low-resource settings. It also leads to wastage when vaccines must be discarded after they have been (potentially) exposed to heat or freezing, which is the main cost driver, adding to the costs of the cold chain itself. There might even be a risk that vaccines with reduced potency are administered, leaving people vulnerable to disease. Alternatively, to facilitate local vaccine production, BioNTech has planned to ship vaccine production units called “BioNTainers” to Africa and other regions left behind during the pandemic. However, this approach is costly and requires a complex implementation.

Unlike mRNA, DNA is considered to be thermostable and does not need the cold chain for transport and storage in environments and countries with limited resources, an essential feature that affects the overall cost of these treatments. The cost of cold chain for DNA stability is an issue that has also been addressed in the context of long-term information storage [9,11]. The research on analytical methods of DNA integrity in different storage conditions has produced a series of methods to evaluate DNA integrity. The same methods are used in gene therapy and genetic vaccination approaches. Pharmacopeia requires DNA integrity above 80%, defined as the percentage of supercoiled plasmid DNA.

In this study, we show that the plasmid DNA coronavirus vaccine COVID-eVax delivered by EP can be stored for 20 days at room temperature, ranging from 23 °C to 45 °C, maintaining biological functions in vitro such as gene expression, and in vivo, such as induced immune responses. The accelerated stability protocol produced the expected increase in DNA nicking (Figure 1). The DNA kept for 20 days at 45 °C showed similar efficacy results in terms of T cells and antibody responses to those of the GMP-like reference DNA (Figure 2). Although the number of mice responding to the vaccination was reduced, the level of the immune responses was not affected, suggesting that it is more a technical variability. In line with this, the experiment with lyophilized DNA showed a more uniform result (Figure 3). Indeed, EP technology is prone to a certain level of variability, as we have observed in murine cancer models [5]. Moreover, it is worth mentioning the indirect evidence observed with linear DNA produced by PCR, which is as immunogenic and efficacious as supercoiled plasmid DNA [12]. In this recent paper, we compared side-by-side PCR-produced linear DNA and plasmid DNA in gene expression and a cancer vaccine model, and showed that the two forms of DNA vaccine delivered by EP resulted in similar immune responses and antitumor effects.

Our results also support the possibility of using COVID-eVax stored at -20 °C for a long time as suggested by the results of DNA kept at 45 °C, which corresponds to storage for 5 years at -20 C° [9]. The limitation of our work is that we did not explore other delivery methods, such as air-mediated jet injection or liponanoparticles with nicked DNA. The possibility of using DNA vaccines without an expensive cold chain may have an economic impact on countries with limited resources for the distribution of pandemic vaccines. Reduced logistic requirements may enlarge the distribution to remote populations and, therefore, may have an impact at the population level.

## Author Contributions

Conceptualization, L.A. and F.P; methodology, F.D., L.L., E.S., A.M., M.C. E. P., and G.B.; validation, G.R. and A.C.; writing—original draft preparation, F.P.; writing—review and editing, L.A. All authors have read and agreed to the published version of the manuscript.

## Institutional Review Board Statement

All the in vivo experimental procedures were approved by the local animal ethics council and the ethical committee of the Italian Ministry of Health, authorization # 586/2019-PR.

## Acknowledgments

We thank Monica Fraternali and Cristina Nardacci for their secretarial assistance.

